# A Draft Male Genome Assembly of the Slipper Lobster *(Thenus australiensis)* Reveals an XY System and a Validated Diagnostic Marker for Monosex Aquaculture

**DOI:** 10.64898/2026.06.24.734161

**Authors:** Ai Hang Tran Nguyen, Gia Huy Ha, Duc Phuc Tran, Nhat Thong Le, Susan Glendinning, Quinn P. Fitzgibbon, Gregory G. Smith, Volker Herzig, Phuc Loi Luu, Tomer Ventura

## Abstract

The slipper lobster (*Thenus australiensis*) is rapidly emerging as a high-potential species for commercial aquaculture. Because females exhibit superior growth characteristics due to less frequent moulting after sexual maturity, developing monosex breeding strategies is highly desirable for industry profitability. However, the lack of genomic resources and early sex-identification tools has hindered this development. Here, we report the first draft male genome assembly for *T. australiensis*, generated using a combination of whole-genome shotgun sequencing, DArT-seq, and multi-tissue transcriptomics. The curated assembly spans 0.913 Gbp with high functional completeness (93.0% BUSCO), providing a robust repertoire of 30,100 protein-coding genes. Through k-mer subtraction and population-level DArT-seq genotyping, we provide definitive evidence that *T. australiensis* utilizes an XX/XY sex-determination system. Crucially, by identifying male-specific structural variations within a neo-Y locus, we developed a diagnostic PCR assay targeting a male-exclusive sequence. This 171 bp marker achieved 100% accuracy in phenotypic sex identification across wild-caught populations. Ultimately, these foundational genomic resources, combined with a highly reliable molecular sexing tool, provide the critical framework necessary for early sex sorting, broodstock management, and the commercial advancement of monosex slipper lobster farming.

**Highlights:**

- First draft genome of slipper lobster (0.913 Gbp) assembled.
- Multi-omics approach confirms an XX/XY sex-determination system.
- Developed a 171 bp diagnostic PCR sex marker with 100% accuracy.
- Facilitates early sex sorting for profitable monosex aquaculture.

## 1. Introduction

*Thenus* spp. are significant seafood species commonly found in the Indo-West Pacific region. In Australia, however, the availability of *Thenus* spp. is primarily restricted to bycatch from scallop and shrimp fisheries, with only a limited supply from industrial hatcheries (Williamson, 2023). Among these, the slipper lobster (*Thenus australiensis*, SL) is increasingly recognized as a promising candidate for sustainable onshore aquaculture development in Australia (Tran Nguyen et al., 2025) due to its shortened larval phase (3–4 weeks compared with 6 months in *Panulirus ornatus*) and its high growth rate (Vijayakumaran and Radhakrishnan, 2011). Interestingly, females tend to be larger than males in SL, as males molt less frequently after reaching sexual maturity (L. Lavalli et al., 2020). It is therefore likely that a monosex population of this species would lead to faster growth, which is expected to improve aquaculture output.

Understanding the biological regulatory pathways involved in sex determination and sexual differentiation is vital for effective monosex breeding strategies in crustaceans. This knowledge also lays a foundation for developing biotechnological tools that enable the production of monosex populations (Tran Nguyen et al., 2025; Wahl et al., 2023). The genetic code forms the foundation of life, and decoding genome sequences provides a valuable way to gain new insights into the molecular processes behind sex determination and differentiation in crustaceans, offering essential tools to address the challenges in aquaculture (The Aquaculture Genomics, Genetics and Breeding Workshop et al., 2017, p. 19). Unfortunately, there remains a significant lack of genomic resources for SL, resulting in a notable gap in both fundamental biological research and practical aquaculture applications, such as biotechnologies for sex manipulation. In implementing these biotechnologies, the availability of sex-specific markers is a vital tool that serves as a valuable genetic resource for examining sex-determining mechanisms and sexual manipulation (Griffiths and Tiwari, 1993). Additionally, it has the potential to significantly reduce the time required for progeny testing, which is typically labor-intensive and time-consuming within the proposed biotechnological framework (Ventura et al., 2011). Various molecular techniques have been explored for sex-marker identification in aquatic species, including whole-genome shotgun sequencing (WGS) combined with genomic analysis (Yang et al., 2020; Zhu et al., 2024); restriction site-associated DNA sequencing (RAD-seq) (Lambert et al., 2016; Shi et al., 2018); type IIB endonuclease restriction-site associated DNA sequencing (2b-RAD-seq) (Cui et al., 2021); randomly amplified polymorphic DNA (RAPD) (Vale et al., 2014); and Diversity Arrays Technology sequencing (DArTseq) (Whankaew et al., 2024). Among these, DArTseq is a hybridization-based technique that offers an affordable, high-throughput system for genetic marker analysis and requires only a small amount of DNA. This reliable method can provide extensive genome coverage, even for organisms without any available reference genome (Jaccoud, 2001). However, DArTseq has specific limitations that must be considered. It targets genomic regions near specific restriction enzyme cut sites, meaning it does not cover all parts of the genome (Alam et al., 2018). Such regions may be underrepresented or completely missing in DArTseq libraries due to uneven genome coverage or bias introduced by restriction digestion, potentially leading to the omission of key sex-linked markers (Alam et al., 2018). In addition, the absence of a reference genome for SL makes genetic sex-marker investigation highly challenging when relying on DArTseq alone.

To overcome the limitations of using reduced-representation methods like DArTseq in an unsequenced organism, whole-genome sequencing is required. Since the introduction of third-generation sequencing technologies in 2011, the application of long-read platforms such as Pacific Biosciences (PacBio) and Oxford Nanopore Technologies has significantly enhanced the generation of high-quality, contiguous genome assemblies (Espinosa et al., 2024; Lee et al., 2016). Despite these advantages, the associated costs of long-read sequencing often limit its widespread application. As a result, second-generation sequencing (short-read techniques), such as WGS, offers a viable alternative for achieving comprehensive genomic insights to support fundamental research. For example, the Illumina platform remains the most widely used technology because of its broad utility, cost-effectiveness, and high accuracy (error rate < 0.1%) (Glenn, 2011). However, *de novo* genome assemblies generated exclusively from short reads can exhibit significant deficiencies, including fragmented assemblies, missing core genes (Li et al., 2010), and poor resolution of repetitive elements (Baptista et al., 2018). For these reasons, it is crucial to implement a multi-platform approach that integrates diverse sequencing methods, analytical techniques, and software. This strategy enables the comprehensive capture of complex genomic regions while minimizing the biases associated with any single method. In this context, the remarkable advancement of transcriptomics (RNA-seq) has provided a powerful tool for exploring molecular pathways, detecting differential gene expression (Chandhini and Rejish Kumar, 2019), and supporting the *de novo* assembly and annotation of high-quality genomes in aquaculture (Kawato et al., 2021). In our previous study on SL, we generated a comprehensive transcriptomic dataset from 6 males and 6 females (40–50 g), with total RNA isolated from 38 samples representing multiple tissues, including the eyestalk, brain, gonads, hepatopancreas, heart, stomach, walking legs associated with the gonopores, and tail muscle. Each sample was sequenced to a depth of at least 20 million reads (Banks et al., 2026). This massive, multi-tissue transcriptomic resource provides strong transcriptional evidence and is a highly valuable dataset for guiding genome assembly and structural gene annotation.

In this study, by integrating short-read platforms (WGS and DArTseq) with multi-tissue RNA-seq data, we report the first draft male genome assembly of SL. Following rigorous decontamination, the final curated genome spans 0.913 Gbp with a high functional BUSCO completeness of 93.0%. We provide strong genomic and transcriptomic evidence that SL utilizes an XX/XY sex-determination system, characterized by males carrying heterozygous, Y-linked regions. Translating this discovery into application, primers designed from a male-specific insertion successfully amplified a 171 bp PCR product exclusively in males, achieving 100% accuracy in phenotypic sex identification. These findings not only offer a highly reliable diagnostic tool for early sex identification but also elucidate the foundational XX/XY system, providing the crucial biotechnological framework needed to develop profitable monosex aquaculture techniques in slipper lobster farming.

## 2. Materials and Methods

### 2.1. Sample collection

Fifteen mature males and fifteen mature females were purchased from fisherman, who collected the animals from Tin Can Bay, Queensland, Australia, for Whole Genome Shotgun, DArTseq, and sex-marker validation. Muscle tissue from the swimming legs was collected from each individual, immediately immersed in 70% ethanol, and stored at −20 °C for further processing. Thirty samples (15 samples/gender) were sent to Diversity Arrays Technology (DArT) Ltd. (DArT Ltd, Building 3, Level D, Kirinari Street, University of Canberra, Bruce, ACT Australia 2617) for DArTseq. Six samples (3 samples/gender) were sent to Australian Genome Research Facility Ltd. (AGRF Ltd, Gehrmann Laboratories, Brisbane QLD, Australia) for DNA extraction and WGS sequencing.

### 2.2. DArT-seq DNA extraction, genotyping and filtering the raw data

DNA was extracted from SL walking legs tissue samples using the NucleoMag kit. (MACHEREY-NAGEL). For the lysis step, samples were overlaid with 50µL of T1 Buffer and 6.25µL of proteinase K. The plate was centrifuged briefly (30-60 sec at 1000 rpm) to ensure the tissue sample was completely submerged in the solution. Samples were then digested overnight at 60 °C, before being centrifuged for 10 min at 3000 rpm and the clear lysate transferred to a new deep well plate. DNA was bound to NucleoMag B-beads using a suspension of 6µL beads in 90µL MB2. The plates were continuously agitated to prevent the beads from settling. Samples were then transferred to the Tecan T100 robot (T100), with the final extraction steps (washing and elution into Elution Buffer) performed on the T100 using 96 tips head.

We received 30 samples, but the DNA quality of 3 samples did not meet sequencing requirements, so we repeated 1 male and 9 females, for a total of 37 assays. Therefore, we obtained 37 assays of 27 samples, processed as follows: Genomic DNA was first incubated with the *PstI* and *SphI* restriction enzymes for digestion to generate fragments with 5′ overhangs. After digestion, samples were cleaned using magnetic beads to remove enzymes and short fragments. Adaptor ligation (adaptor sequences are the proprietary of DartSeq company) was then performed using enzyme-specific adaptors compatible with the *PstI* and *SphI* cohesive ends in the presence of T4 DNA ligase. Each adaptor pair contained a flow-cell attachment sequence, sequencing primer binding site, and a sample specific barcode. Ligation products were PCR-amplified to selectively enrich mixed end fragments and to complete Illumina sequencing constructs^27^. The *PstI* and *SphI* compatible adaptors were designed to include the Illumina flowcell attachment sequence, sequencing primer sequence and “staggered” which consists of varying length barcode region, similar to the sequence reported by (Elshire et al., 2011). The reverse adaptor contained the flowcell attachment region and SphI compatible overhang sequence. Only “mixed fragments” (i.e., fragments flanked by *PstI* and *SphI* ends) were effectively amplified in 30 rounds of PCR using the following reaction conditions: 1. 94 °C for 1 min 2. 30 cycles of: 94 °C for 20 sec 58 °C for 30 sec 72 °C for 45 sec 3. 72 °C for 7 min After PCR, equimolar amounts of amplified product were bulked and subjected to 100 cycles of sequencing (single reads) on the Illumina NovaSeq X+ sequencer. Sequences generated from each lane were processed using proprietary DArT analytical pipelines. In the initial pipeline, poor-quality sequences were removed, with more stringent filtering applied to the barcode region than to the rest of the sequence, ensuring reliable assignment of sequences to specific samples during barcode demultiplexing.

Following filtration (Barcode region Min Phred pass score 30, Min pass percentage 75, Whole read Min Phred pass score 10, Min pass percentage 50), approximately 238.633 unique sequences per sample were used in marker calling. Identical sequences were collapsed into “fast call files”, which were “groomed” using DArT PL’s proprietary algorithm (a script written in the R programming language specifically designed to analyse genetic marker data generated by DArT). This algorithm corrects low-quality bases from singleton tags into correct bases using collapsed tags with multiple members as a template. The “groomed” fastqcol files were then used in the secondary pipeline for DArT PL’s proprietary SNP and SilicoDArT (presence/absence of restriction fragments in representation) calling algorithms (DArTsoft14).

### 2.3. AGRF DNA extraction, library preparation and whole genome shotgun sequencing

Genomic DNA was extracted using the Qiagen DNeasy Plant Mini kits (Qiagen Pty Ltd, 1894 Dandenong Road, Clayton VIC 3168, Australia) following the manufacturer’s instructions. 200 ng - 300 ng/sample used as input DNA for WGS. Library preparation was performed using the Illumina® DNA PCR-Free Prep, Tagmentation (96 Samples) kit (Cat# 20041795), following the manufacturer’s instructions for all steps. Sequencing was done using paired-end 150 bp reads on Illumina’s NovaSeq X Plus system with RTA version 4.6.7.

### 2.4. Genome Investigation

2.4.1. **Genome Size Estimation**

The sequencing data used in this study were generated using two library preparation methods and two sequencing platforms, providing a comprehensive dataset for genome investigation. The Whole-Genome Sequencing (WGS) library, prepared from 3 male and 3 female samples, was sequenced on the Illumina NovaSeq X+ platform, producing the most extensive dataset (380.00 Gbp) with 150-bp reads and the highest coverage (300X) (see Supplementary Table S1). The Diversity Arrays Technology Sequencing (DaRT-Seq) library, from 15 male and 22 female samples on the Illumina NovaSeq X+ platform, provided a smaller targeted dataset of 7.56 Gbp with 120 bp reads, achieving a lower average coverage of 6X. Finally, the RNA-Seq data were obtained from a previous study (Banks et al., 2026) (PRJNA772049) available at NCBI, totalling 290.78 Gbp and averaging 150 bp, yielding 232X coverage (see Supplementary Table S1).

We processed six Whole Genome Sequencing (WGS) and 27 DArT-seq samples to ensure high-quality data. Quality control was performed using fastp (v0.23.4) by trimming 8 bp from the 5′ end of all reads, filtering out bases with a Phred quality score below 25, and discarding reads shorter than 50 bp. Following this initial cleaning, the WGS reads were pooled by sex into single male and female datasets. To estimate genome size and heterozygosity, we performed k-mer analysis using KMC (v3.2.1). We generated k-mer frequency histograms for a range of k-values (k=18 to 54 with a step size of 3) by counting canonical k-mers in the merged paired-end reads (R1 and R2 counted separately and then united). The resulting histograms were modeled using GenomeScope (v2.0) (Ranallo-Benavidez et al., 2020; Vurture et al., 2017).

#### 2.4.2. Genome Assembly

*De novo* assembly was performed using pooled male WGS reads. We first normalized the read coverage to a target depth of 40× using bbnorm (v39.33) (Bushnell, 2014) to reduce data redundancy and complexity. These normalized reads were then assembled using ABySS (v2.3.10) (Jackman et al., 2017) with a k-mer size of 65. To improve the continuity of the assembly, the draft genome was scaffolded using short-read RNA-seq alignments with Rascaf (v20180710) (Song et al., 2016), followed by a long-range scaffolding step using L_RNA_scaffolder (https://github.com/CAFS-bioinformatics/L_RNA_scaffolder) and our *de novo* assembled transcripts. The scaffolded assembly was subsequently polished for three iterative rounds using the raw short-read DNA-seq data and ntEdit (v1.4.3) (Warren et al., 2019) with k-mer sizes of 31, 41, and 25 to correct base errors and indels.

#### 2.4.3. Genome Curation and Quality Control

Following the initial *de novo* assembly and polishing steps, the draft genome was subjected to rigorous quality control to identify and remove foreign DNA and environmental contamination. We utilized the National Center for Biotechnology Information (NCBI) Foreign Contamination Screen (FCS-GX) tool (Astashyn et al., 2024), which evaluates assembly sequences against a comprehensive database of known contaminants—including bacteria, fungi, and viruses—to detect non-target genomic elements. Contigs definitively flagged by FCS-GX as putative environmental or microbial contamination, as well as highly fragmented scaffolds lacking robust read support, were curated and permanently excluded from the dataset. This stringent decontamination process refined the assembly down to a final, polished male genome of 0.913 Gbp. All subsequent downstream analyses, including structural gene prediction, functional annotation, and repeat masking, were executed exclusively on this curated genomic dataset to ensure that the resulting biological inferences accurately reflected the *T. australiensis* genome without confounding microbial noise.

#### 2.4.4. Transcriptome Assembly

A *de novo* transcriptome was generated to support genome annotation using the nf-core/denovotranscript pipeline (v1.2.1) (Ewels et al., 2020), which employs the Trinity assembler (Haas et al., 2013). To create a high-confidence, non-redundant dataset, we applied a multi-step filtering process. Transcripts were first clustered at 97% sequence identity using CD-HIT-EST (v4.8.1) (Fu et al., 2012), after which expression levels were quantified using Salmon (v1.10.3) (Patro et al., 2017). Transcripts with low expression (TPM < 1) were removed, and a final clustering step was performed at 99.5% identity to eliminate remaining redundancy while preserving sequence variation.

#### 2.4.5. Annotation of Repeat Regions

We employed a combination of *de novo* and homology-based methods to identify repetitive elements within the genome. Repeat families were first identified *de novo* using RepeatModeler (v2.0.1) (Flynn et al., 2020). We then used RepeatMasker (v4.1.2) (accessed on 1/7/2025 https://www.repeatmasker.org/RepeatMasker/) (Smit et al., 2013) to screen the genome against these *de novo* repeats and the Repbase library. Full-length long terminal repeats (LTRs) were identified using LTR_Finder (v1.07) (Xu and Wang, 2007), while simple sequence repeat (SSR) markers were detected using MISA (v2.1) (Beier et al., 2017).

#### 2.4.6. Gene Structure Prediction and Annotation

For structural annotation, we selected contigs larger than 2 kb and utilized a combined approach integrating ab initio gene prediction with transcript evidence. Gene structures were predicted using AUGUSTUS (v3.5.0) (Stanke et al., 2006) via the BRAKER (v1.9) (Hoff et al., 2019) pipeline, which used RNA-seq alignments as training evidence, while simultaneously aligning transcripts to the masked genome using PASA (v2.5.3) (Haas, 2003). These independent predictions were merged into a single weighted consensus gene set using EVidenceModeler (EVM v2.5.3) (Haas et al., 2008), and the consensus models were subsequently updated using PASA to refine untranslated regions (UTRs). Functional annotation was achieved by aligning predicted protein sequences against the NCBI Non-Redundant (NR - downloaded 2025-10-11) and Swiss-Prot (downloaded 2025-10-11) databases using DIAMOND (v2.1.8) (e-value cutoff 10^-5^) (Buchfink et al., 2021, 2021), with protein domains, GO terms, and orthology assigned using InterProScan (v5.52) (Jones et al., 2014) and eggNOG-Mapper (v2.1.9) (Huerta-Cepas et al., 2017).

#### 2.4.7. Non-coding RNA Detection

The genome was screened for non-coding RNAs (ncRNAs) using a suite of specialized tools. Transfer RNAs (tRNAs) were predicted using tRNAscan-SE (v2.0.12) (Chan et al., 2021), while ribosomal RNAs (rRNAs), microRNAs (miRNAs), and small nuclear RNAs (snRNAs) were identified by searching the Rfam database (v14.5) (Kalvari et al., 2021) using Infernal (v1.1.4) (Nawrocki and Eddy, 2013). Additionally, small nucleolar RNAs (snoRNAs) were predicted using Snoscan (v1.0) (Lowe and Eddy, 1999), utilizing the identified rRNAs as target sequences.

#### 2.4.8. Phylogenetic Analysis

To elucidate the evolutionary relationship of SL within the subphylum Crustacea, we compiled a dataset of protein sequences from 39 reference species available in the NCBI database. Orthologous sequences were identified using BLASTp (v2.13.0) (Camacho et al., 2009), after which the best-scoring hits for each ortholog were concatenated to form a supermatrix. We performed multiple sequence alignment using MAFFT (v7.511) (Katoh, 2002) to ensure accurate residue correspondence. Subsequently, a phylogenetic tree was constructed using IQ-TREE (v1.6.12) (Nguyen et al., 2015)under the BLOSUM62 substitution model (Henikoff and Henikoff, 1992). The final phylogenetic reconstruction was visualized using the ggtree package (Yu et al., 2018).

### 2.5. Sex-marker Investigation

2.5.1. **Acquisition of the Candidate Sex-specific Markers and Positive Controls**

Sex-specific markers: To identify candidate sex-specific DNA markers, we employed a k-mer subtraction and assembly strategy utilizing the ssp2 pipeline (https://github.com/fengtong-bio/ssp2) (Ruan et al., 2021). First, canonical k-mer databases (k=21, 33, 55) were generated for both the WGS and DArT-seq datasets using KMC (v3.2.1) (Kokot et al., 2017). We then computationally isolated sex-specific k-mers by subtracting the male k-mer pool from the female datasets, and vice versa. Reads containing these unique k-mers were extracted using BBDuk (v39.33) (Bushnell, 2014) and assembled de novo using MEGAHIT (v1.2.9) (Li et al., 2015) with an iterative k-mer list of 27–141 to generate candidate sex-specific contigs. To validate these candidates, WGS and DArT-seq reads from multiple male and female individuals were mapped back to the contigs using BWA-MEM (v0.7.17-r1188) (Li and Durbin, 2009). Per-base read depth was calculated using SAMtools (v1.3.1) (Danecek et al., 2021), and custom filtering scripts were applied to identify sequences exhibiting strict sex-biased coverage—specifically isolating markers with high coverage in one sex and an absolute absence of coverage in the other. Positive controls: To ensure the reliability of our mapping and sequencing pipeline, we identified “positive control” markers—sequences that must be present in all individuals. Instead of filtering for sex bias, a custom AWK script was used to process the read depth data. This script filtered for genomic positions where every individual sample, regardless of sex, had a read depth greater than zero (Depth > 0). These positions were then grouped into continuous blocks, allowing a maximum gap of 10 base pairs (GAP_LENGTH = 10). We only retained regions with a minimum length of 100 base pairs (MIN_LENGTH = 100). This method provided a set of high-confidence, universal markers that validated our pipeline’s ability to consistently detect DNA across all male and female samples.

#### 2.5.2. Genomic Mapping, Coverage Depth Analysis, and Statistical Validation Pipeline

To determine the physical locations and genomic contexts of the candidate male-specific markers, the assembled sex-specific contigs were mapped to the *T. australiensis* whole-genome assembly. Homology searches were performed using the BLASTn algorithm with default parameters and an *E-value* threshold of 10^-5^. The resulting high-confidence alignments were analyzed to identify the specific genomic scaffolds and coordinates harboring the putative Y-linked insertions.To statistically validate the sex-specificity of these candidate regions, raw paired-end and trimmed single-end reads from the male (n = 18) and female (n = 25) cohorts were aligned to the candidate male-specific scaffold (Scaffold_2453) using BWA-MEM (v0.7.17). Alignments were sorted and indexed using SAMtools (v1.9). Sequencing depth was calculated in non-overlapping 1,000 bp windows using Mosdepth (v0.3.10) (Pedersen and Quinlan, 2018). Coverage data were analyzed using a custom Python pipeline employing the pandas and scipy.stats libraries. For each genomic window, the normalized median depth was calculated for both sexes. Male bias was quantified using the log2 ratio of male-to-female depth, calculated as log2((Median_Male + ε) / (Median_Female + ε)). Significance was assessed using a two-sided Mann-Whitney U test. Genomic windows were definitively classified as Y-linked if they exhibited a male median depth > 0.5, a female median depth < 0.05, and a log2 ratio > 3.0.

#### 2.5.3. Functional Annotation of Sex-specific Markers

To determine the biological function of the in silico identified sex-specific markers, the validated genomic contigs were searched against the predicted lobster proteome using BLASTx with a strict E-value threshold of 10^-5^. Top-scoring protein accessions were extracted and cross-referenced with functional data from the InterProScan, NCBI Non-Redundant (NR), Swiss-Prot, and eggNOG-mapper databases to provide an integrated annotation of protein domains, orthologous groups, and curated functional descriptions. This multi-database approach ensured high-confidence identification of the male-specific genomic loci, ultimately characterizing the functional annotation of the sex-specific markers.

To classify the host gene and characterize the disruptive elements, domain analyses were conducted using PANTHER classification (to identify the *CENP-E* kinesin-7 motor protein). Concurrently, the male-specific insertions residing within the intronic regions were categorized by identifying transposase and reverse transcriptase domains indicative of Class II DNA transposons (Tc1-like) and degraded Class I retrotransposons (LINE-1/Jerky). This multi-database approach ensured a high-confidence characterization of the transposon-mediated pseudogenization occurring at the neo-Y locus.

#### 2.5.4. Transcriptomic profiling and structural analysis of the neo-Y gene

To elucidate the regulatory isolation and expression divergence of the male-specific loci, we cross-referenced the newly identified neo-Y candidate gene (the truncated *CENP-E* paralog) and its autosomal ancestor against the multi-tissue *T. australiensis* transcriptome atlas (Tran, et al., manuscript in preparation). Transcript variants, specifically the testis-exclusive transcript (NonamEVm002966t3) and the fully functional ancestral transcript (NonamEVm000093t1), were mapped back to their respective host scaffolds (Scc_2453 and Scc_1163) using minimap2. This alignment was utilized to assess the loss of synteny, genomic relocation, and the exact physical breakpoint of the transposon-mediated truncation.

Furthermore, comparative expression profiles spanning comprehensive somatic and reproductive tissues were extracted from the transcriptome database. Expression levels, quantified as Trimmed Mean of M-values (TMM), were analyzed to statistically confirm the strictly testis-exclusive transcription of the neo-Y locus, providing transcriptomic validation of its divergence from the pleiotropically expressed autosomal ancestor.

#### 2.5.5. Validation of Sex-specific Markers Using *In Silico* and Experimental Approaches

To validate the sex-linkage of the candidate genes *in silico*, a comparative read-depth analysis was conducted using raw WGS and DArT-Seq data from 25 female and 18 male samples (3 males and 3 females for WGS and 15 males and 22 females for DArT-Seq, collected from fishermen) . The raw reads were mapped to the de novo assembly, and the resulting alignments were grouped by sex to generate cumulative coverage profiles. Average sequencing depth for the candidate scaffold was calculated by extracting per-base depth values, including positions with zero coverage, to ensure an unbiased quantitative comparison.

To experimentally validate the sex linkage of the candidate genes, SL swimming leg samples from 15 mature males and 15 mature females stored in 70% ethanol were used. Genomic DNA was extracted using the REDExtract-N-Amp Tissue PCR Kit (Sigma, St Louis, MI, USA) Genomic DNA was extracted using the REDExtract-N-Amp Tissue PCR Kit (Sigma, St Louis, MI, USA) according to the manufacturer’s instructions. The Primer3web version 4.1.0 software (Untergasser et al., 2012) was used to design primers for the sex-specific marker. The forward primer was selected within the specific sequence, and the Reverse primer was outside the sequence (see Supplementary Table S7.1), called SL-SM1. SL-PC sequences are used as a positive control. PCR conditions included preheating to 94 °C for 3 min, followed by 30 cycles of 94 °C for 30 s, 59 °C for 30 s, and 72 °C for 2 min, followed by final elongation at 70 °C for 10 min and a final hold at 4 °C. PCR products were electrophoresed on a 2% agarose gel and visualized under UV light with ethidium bromide. All the primers are listed in Table 1.

**Table 1.**
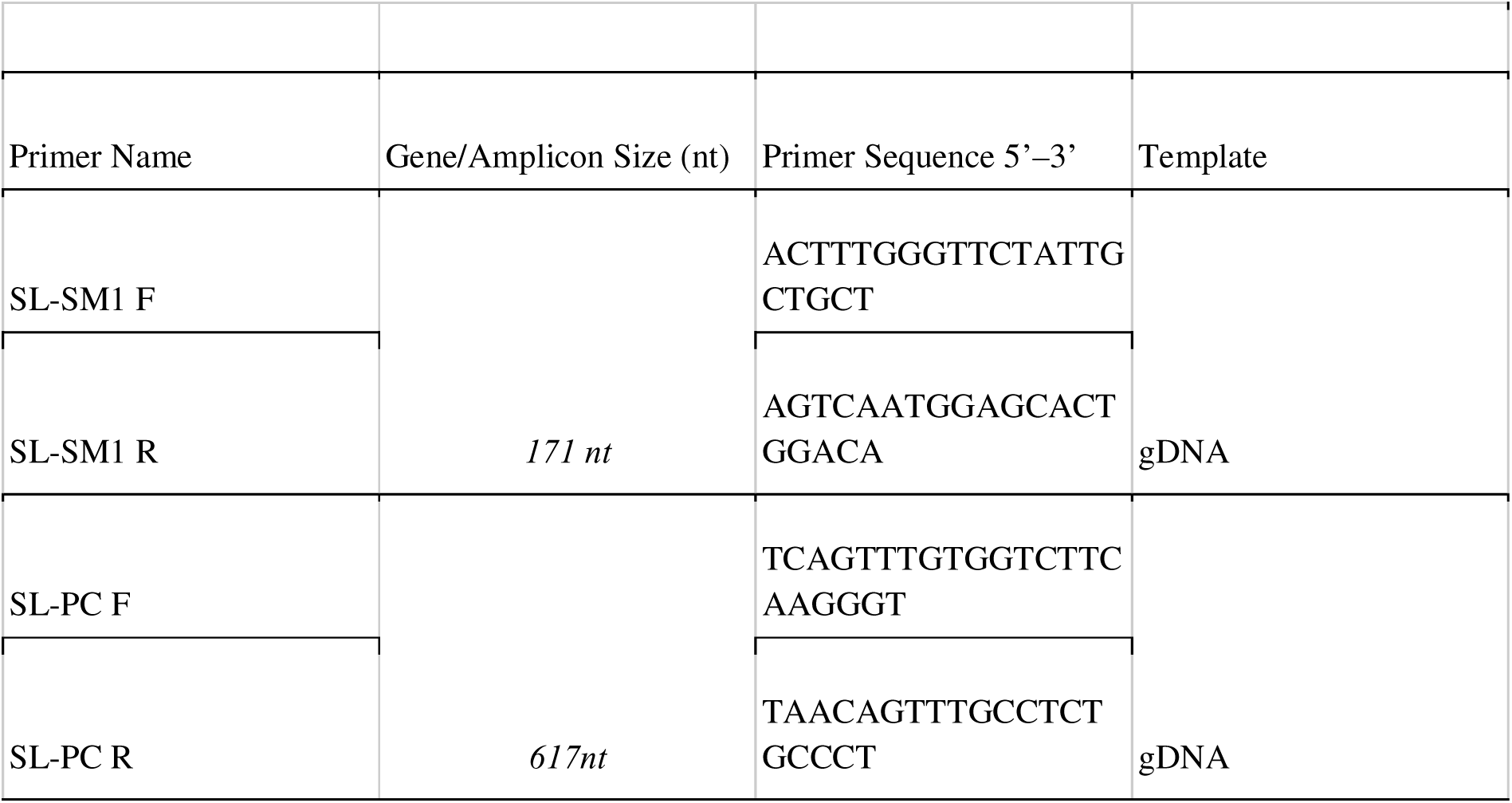
Male-specific marker primers.

### 2.6. Discovery of the sex chromosome system

DArT-Seq results were used to analyse the sex chromosome system in SL. Following genomic DNA extraction and library construction, a total of 14 male and 13 female samples were sequenced and analysed. Male or female-biased alleles were investigated, with the ratio of males to females carrying the allele at least 1:2. Using ClustVis, a heatmap was generated showing clear sex segregation between the male and female samples. Then, male-specific sequences were downloaded as a fasta file and uploaded to CLC (v.8) and BLASTed against the SL transcriptome library using the male reproductive-related RNA-Seq files (3 x testis samples + 5th walking leg sample) as well as the female reproductive-related RNA-Seq (3 x ovary samples + 3rd walking leg sample) to confirm SNP bias.

## 3. Results

### 3.1. Genome Investigation

#### 3.1.1. Genome size estimate

We conducted genome surveys to estimate key characteristics of the male and female *Thenus australiensis* (SL) genomes, including overall size, heterozygosity, and repeat content. Using GenomeScope, we evaluated K-mer sizes ranging from 18 to 54. At the optimal K-mer size of 33, we observed dominant peak depths of 30.6 for females and 27.2 for males. Based on these distributions, the estimated genome size of males (∼1.25 Gbp) was slightly larger than that of females (∼1.12 Gbp) (Figure 1A, 1B).

**Figure 1.**
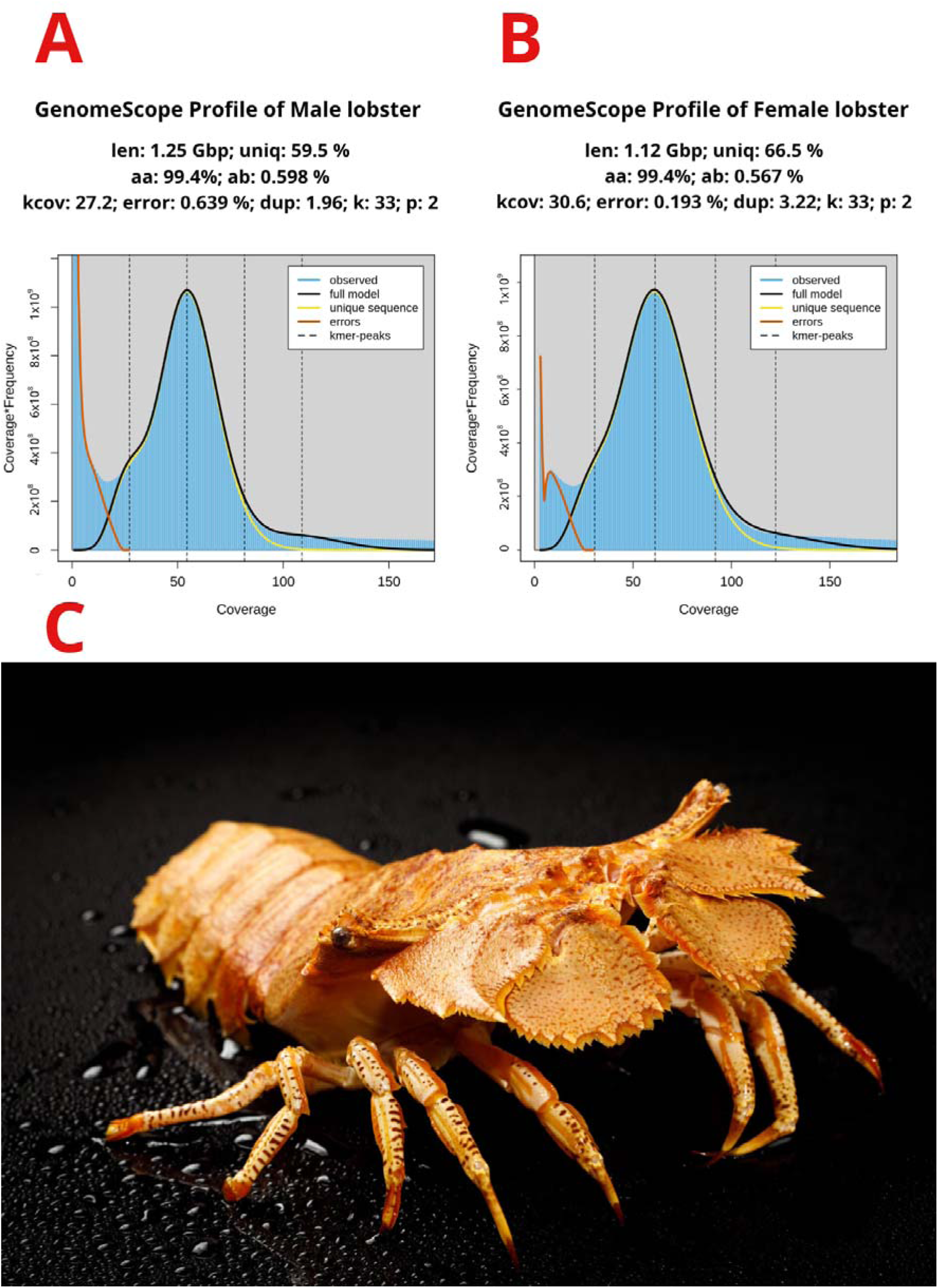
Genome size estimation and morphological characteristics of the male and female slipper lobster (*Thenus australiensis*). **A)** Genome size estimation of male SL with k=33; **B)** Genome size estimation of female SL with k=33. Genomescope report—coverage of the kmer (x) by kmer counts (y) with k-mer = 33 with Len, estimated total genome length; Uniq, unique portion of the genome (not repetitive); Het, heterozygosity rate; Kcov, K-mer coverage for the heterozygous bases; Err, error rate; Dup, duplication rate, where the blue bar in the graph represents the observed K-mer, and the yellow and orange liines in the graph represent the errors and unique sequences, respectively—on the left, the coverage represented is the linear plot, and on the right, the coverage represented is log-scale; **C)** *Thenus australiensis* picture.

Males also exhibited a marginally higher heterozygosity rate (0.59%) compared to females (0.56%). Unique sequence content accounted for 59.5% of the male genome and 65.5% of the female genome, indicating a higher proportion of repetitive sequences in males. Notably, the sequencing error rate was significantly higher in the female dataset (0.639%) than in the male dataset (0.193%). This disparity in data quality led us to prioritize the male dataset for the subsequent de novo assembly to ensure the most accurate and comprehensive results. Overall, the k-mer frequency distributions for both sexes were consistent with a diploid genome model, characterized by a single prominent peak for unique sequences and an extended tail representing repetitive regions (Supplementary Table S2).

#### 3.1.2. Male genome sequencing, curation, and assembly

Our initial de novo sequencing effort produced a preliminary draft male genome spanning 1.44 Gbp. Because this initial assembly was significantly larger than our k-mer-based estimate of 1.25 Gbp, we suspected the presence of environmental or microbial DNA. To ensure data integrity, the draft assembly was rigorously screened using the NCBI Foreign Contamination Screen (FCS) tool. We permanently removed any contigs definitively flagged as bacterial or environmental contamination, alongside highly fragmented sequences lacking robust read support.

This essential curation step yielded a final, polished genome assembly of 0.913 Gbp with a GC content of 41.79% (Table 2). While the genome remains highly fragmented—a common characteristic of assemblies relying predominantly on short-read sequencing—the decontamination process dramatically improved its overall structural reliability. The final assembly consists of 808,063 scaffolds (down from over 3.9 million in the uncurated draft) with a scaffold N50 of 3,392 bp and a maximum contig length of 232,493 bp.

**Table 2.** Genome assembly statistics of the male Thenus australiensis.

| Table 2. Genome assembly statistics of the male *Thenus australiensis* |  |
| --- | --- |
| Estimate genome size | 1.25 Gbp |
| Total length | 0.913 Gbp |
| GC content | 41.79 % |
| Largest contig | 232,493 bp |
| Scaffold N50 | 3392 bp |
| Number of scaffolds | 808,063 |
| Number of protein-coding genes | 36,907 |
| Average gene length | 4,214 bp |
| Average CDS length | 1,159bp |

Despite the physical fragmentation, downstream annotation of this clean gene space was highly successful. We predicted a robust set of 36,907 protein-coding genes, exhibiting an average gene length of 4,214 bp and an average coding sequence (CDS) length of 1,159 bp. To evaluate the functional completeness of the curated genome, we performed a BUSCO analysis against the arthropoda_odb12 dataset (Table 3). The assessment confirmed excellent gene recovery despite the removal of contaminated contigs, capturing 93.0% (1,550) of highly conserved arthropod orthologs. The vast majority were complete and single-copy (92.5%, 1,542), with only a negligible fraction appearing as duplicates (0.5%, 8). The remaining core genes were classified as either fragmented (5.4%, 90) or missing (1.6%, 27). These metrics confirm that while decontamination reduced the overall physical size of the assembly, the biologically critical coding regions of *T. australiensis* were preserved with high fidelity.

**Table 3.** BUSCO assessment of genome completeness using the arthropoda_odb12 dataset.

|  |  |  |
| --- | --- | --- |
| **Type** | **Number** | **Proportion (%)** |
| **Complete BUSCOs** | 1550 | 93 |
| **Complete and single-copy BUSCOs** | 1542 | 92.5 |
| **Complete and duplicate-copy BUSCOs** | 8 | 0.5 |
| **Fragmented BUSCOs** | 90 | 5.4 |
| **Missing BUSCOs** | 27 | 1.6 |
| **Total** | 1667 | 100 |

#### 3.1.3. Integrated gene structure prediction and functional annotation

Our integrated pipeline identified a refined set of 30,100 predicted gene structures. These robust models displayed an average gene length of 3,298 bp and an average CDS length of 1,129 bp (Supplementary Table S3). A subsequent BUSCO assessment of these predicted models confirmed high-quality annotation, successfully recovering 87.3% of the core arthropod orthologs (77.3% single-copy; 10.0% multi-copy) (Table 4).

**Table 4.** BUSCOs assessment of genome with predicted gene completeness.

| **Table 4. BUSCOs assessment of genome with predicted gene completeness** |  |  |
| --- | --- | --- |
| **Type** | **Number** | **Proportion (%)** |
| **Complete BUSCOs (C)** | 1200 | 72 |
| **Complete and single-copy BUSCOs (S)** | 1089 | 65.3 |
| **Complete and duplicate-copy BUSCOs (D)** | 111 | 6.7 |
| **Fragmented BUSCOs (F)** | 154 | 9.2 |
| **Missing BUSCOs (M)** | 313 | 18.8 |
| **Total BUSCOs** | 1667 | 100 |

Downstream functional analysis further highlighted the comprehensive coverage of the clean genome. Of the 30,420 total predicted genes utilized for mapping, 24,597 were successfully assigned putative biological functions (Supplementary Table S4). The NCBI Non-Redundant (NR) database provided the most extensive mapping, reflecting broad taxonomic conservation. A strong consensus emerged across the analytical platforms, with 8,237 genes confidently annotated across all four major databases: KEGG, Swiss-Prot, InterPro, and NR (Figure 2A).

**Figure 2.**
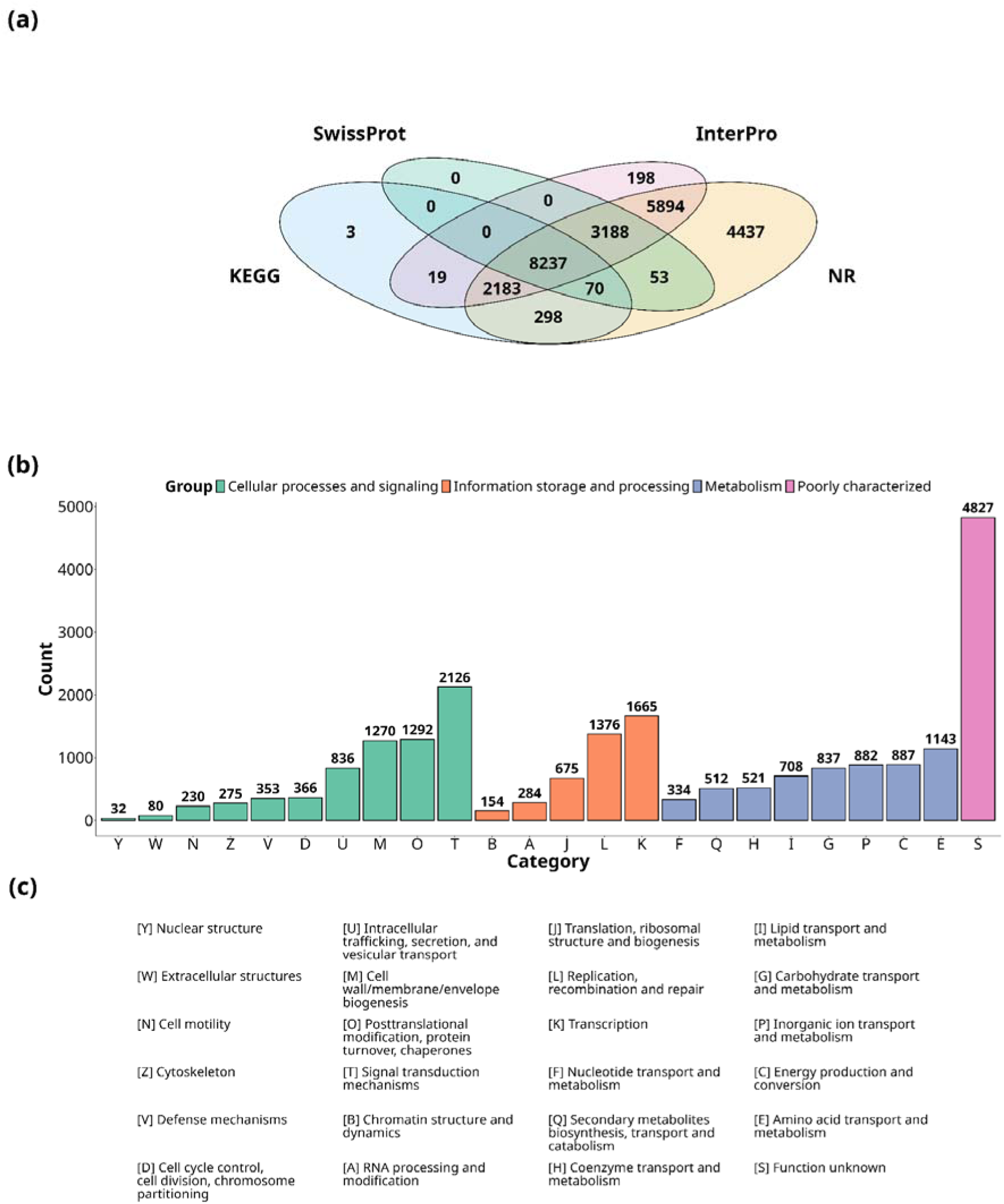
Functional annotation and COG classification of the curated *T. australiensis* gene space. **(a)** Venn diagram illustrating the distribution and overlap of functionally annotated genes across four major databases: KEGG, Swiss-Prot, InterPro, and NCBI Non-Redundant (NR). **(b)** Histogram detailing the classification of predicted genes into Clusters of Orthologous Groups (COG). The x-axis denotes specific COG categories, and the y-axis represents the absolute number of genes assigned to each. Bars are color-coded to reflect broader functional macro-groupings: Cellular processes and signaling (green), Information storage and processing (orange), Metabolism (purple), and Poorly characterized (pink). **(c)** Legend defining the specific biological functions corresponding to each COG category abbreviation presented in panel (b).

Classification via Clusters of Orthologous Groups (COG) revealed that while the largest single category remains “Function unknown” (4,827 genes)—as is expected for a non-model organism—well-defined biological pathways are strongly represented (Figure 2B, 2C). Dominant categories included Signal transduction mechanisms (2,126 genes), Transcription (1,665 genes), Replication, recombination, and repair (1,376 genes), and Amino acid transport and metabolism (1,143 genes). Conversely, highly specialized categories such as Nuclear structure (32 genes) and Extracellular structures (80 genes) were the least represented.

#### 3.1.4. Repeat elements and non-coding RNAs

Owing to the fragmented nature of the short-read genome assembly, a comprehensive annotation of the entire repeatome was not feasible. Nevertheless, simple sequence repeats (SSRs) were successfully identified and classified based on motif length, accounting for 2.23% of the genome (Supplementary Figure S1). Additionally, we successfully identified four distinct types of non-coding RNAs (ncRNAs), cataloging 3,284 snRNAs, 3,036 rRNAs, 2,778 tRNAs, and 388 miRNAs (Supplementary Table S5).

#### 3.1.5. Phylogenetic analysis

To elucidate the evolutionary relationship between SL and other crustaceans, we constructed a phylogenetic tree utilizing 200 identified orthologous genes. The resulting topology demonstrated that non-decapod groups, including Isopoda, Amphipoda, and Stomatopoda, formed distinct outgroup clusters basal to the main lineage (Figure 3). Within the Decapoda clade, SL diverged distinctly from the shrimp lineages, grouping closely with the spiny lobster (*Panulirus ornatus*) and the clawed lobster (*Homarus americanus*). This branching pattern clearly illustrates their evolutionary divergence and robustly positions *T. australiensis* within the lobster clade.

**Figure 3.**
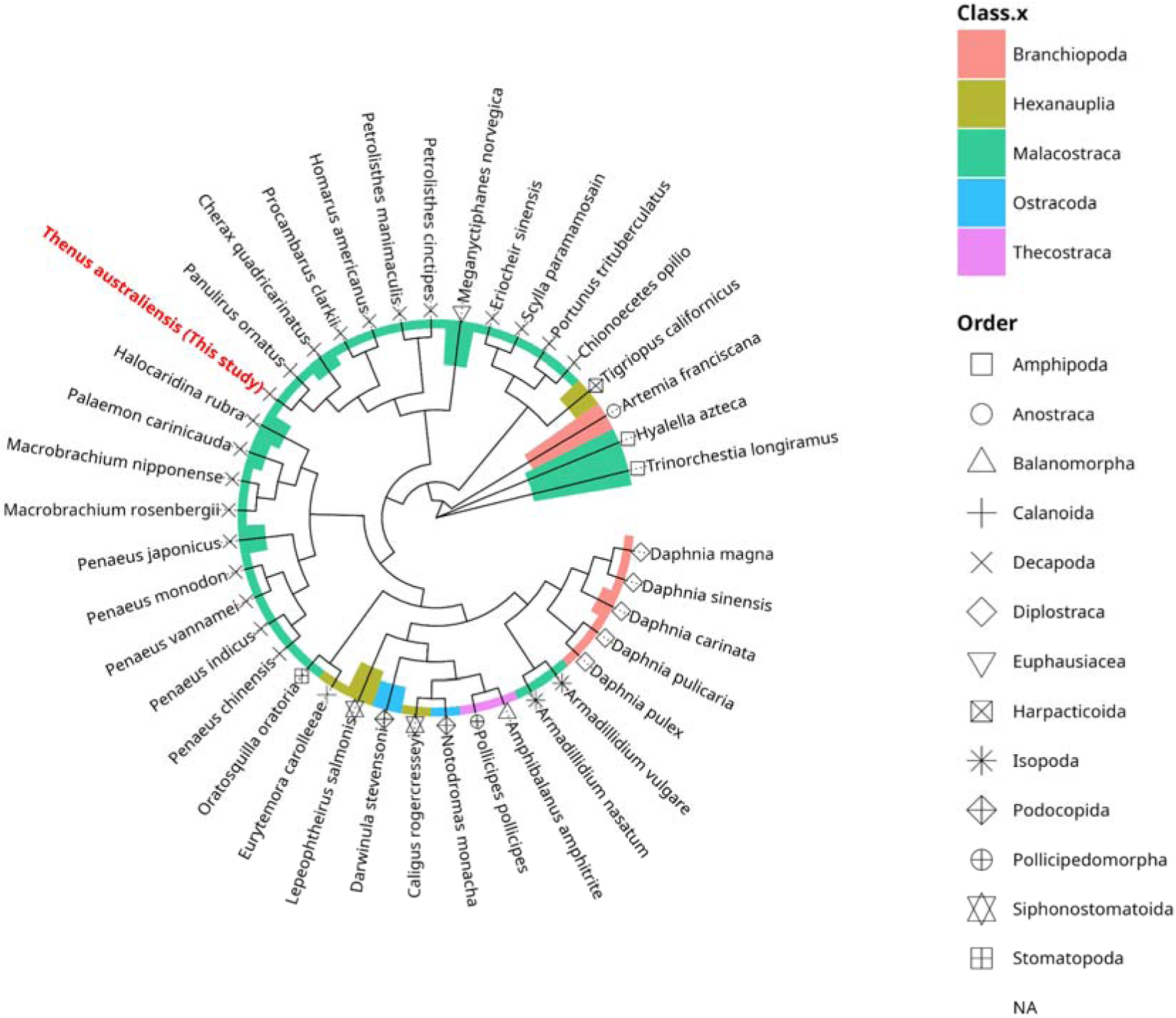
Phylogenetic relationship of *Thenus australiensis* (SL) within the subphylum Crustacea. The maximum likelihood phylogenetic tree was constructed using IQ-TREE based on a concatenated supermatrix of 150 single-copy orthologous genes extracted via BUSCO analysis across SL and 39 reference crustacean species. The inner branch symbols denote specific taxonomic orders, while the outer colored ring classifies the species by taxonomic class. *Thenus australiensis* (highlighted in red text) diverges distinctly from the shrimp lineages, clustering closely with the spiny lobster (*Panulirus ornatus*) and clawed lobster (*Homarus americanus*).

### 3.2. Sex-marker investigation

#### 3.2.1. Identification of candidate sex-specific fragments

The k-mer subtraction and assembly process yielded 951,293,824 and 1,123,647,428 reads localized by female- and male-specific k-mers, respectively. These reads were assembled into 1,684,073 putative female-specific and 2,079,158 putative male-specific contigs (Supplementary Table S6). After mapping clean reads from all individuals back to these sequences to verify their sex-specificity, all female-specific candidates failed the validation step. These sequences either matched several male samples or were not consistently present across all female samples, resulting in no retained female-specific markers.

In contrast, we successfully identified six male-specific contigs (Supplementary Table S7.1). Detailed analysis revealed that within these larger contigs, only a small segment of roughly 100 to 120 bp was truly sex-specific. These precise male-exclusive regions were isolated and used as target sequences for designing diagnostic PCR primers (Supplementary Table S8.1).

For internal positive controls, computational analysis identified 5,660 potential contigs that were consistently detected across all individual samples, demonstrating the technical reliability of our sequencing and mapping pipeline. From these, we selected contig k141_1672385 (4,117 bp) for further validation (Supplementary Table S7.2). We developed a primer set targeting a 208 bp region within this contig. BLAST analysis against the male and female genome assemblies confirmed a perfect match, establishing this universally present marker as a robust internal PCR control.

#### 3.2.2. Genomic architecture of the male-specific region

Analysis of Scaffold_2453 (122,615 bp) revealed distinct male-specific genomic segments. The strict male-exclusivity of these regions was demonstrated by the mapping of our previously identified sex-specific markers. Specifically, BLASTn mapping showed that the male-specific contig k141_7345028 aligns with high similarity (>99.5%) directly to a targeted insertion area (Region A). As illustrated in the read depth heatmap (Supplementary Figure S3), the mapping signal for this contig is entirely restricted to males, visually confirming it as a true male-unique insertion. Structural annotation of this host scaffold revealed that this male-specific insertion interrupts a paralog of Centromere Protein E *(CENP-E).* To determine the functional consequences of these transposon invasions, we cross-referenced the neo-Y *CENP-E* locus against a comprehensive multi-tissue *T. australiensis* transcriptome atlas. This analysis conclusively linked Scaffold_2453 to a severely truncated, testis-specific transcript *(NonamEVm002966t3)*, which shares ancestry with a fully intact *CENP-E* gene *(NonamEVm000093t1)* located on an autosomal scaffold. Expression profiling confirmed that the ancestral copy is pleiotropic, exhibiting broad transcription across somatic tissues with significant upregulation in the gonads of both sexes (Figure 5A). In striking contrast, the newly duplicated neo-Y *CENP-E* paralog has been entirely silenced in somatic tissues, with its transcription strictly restricted to the male testis (Figure 5B). Crucially, the male-specific insertion in Region A consists of a Tc1 transposable element that prematurely halts the transcription of the neo-Y locus. This transposon-mediated truncation actively degrades the male-specific transcript, resulting in a severely truncated protein (Figure 4). This structural variation—characterized by the accumulation of mobile elements and subsequent gene degradation—is a classic genomic signature of an emerging Y chromosome. More importantly, this precise structural breakpoint provided the ideal target for our diagnostic sex marker.

**Figure 4.**
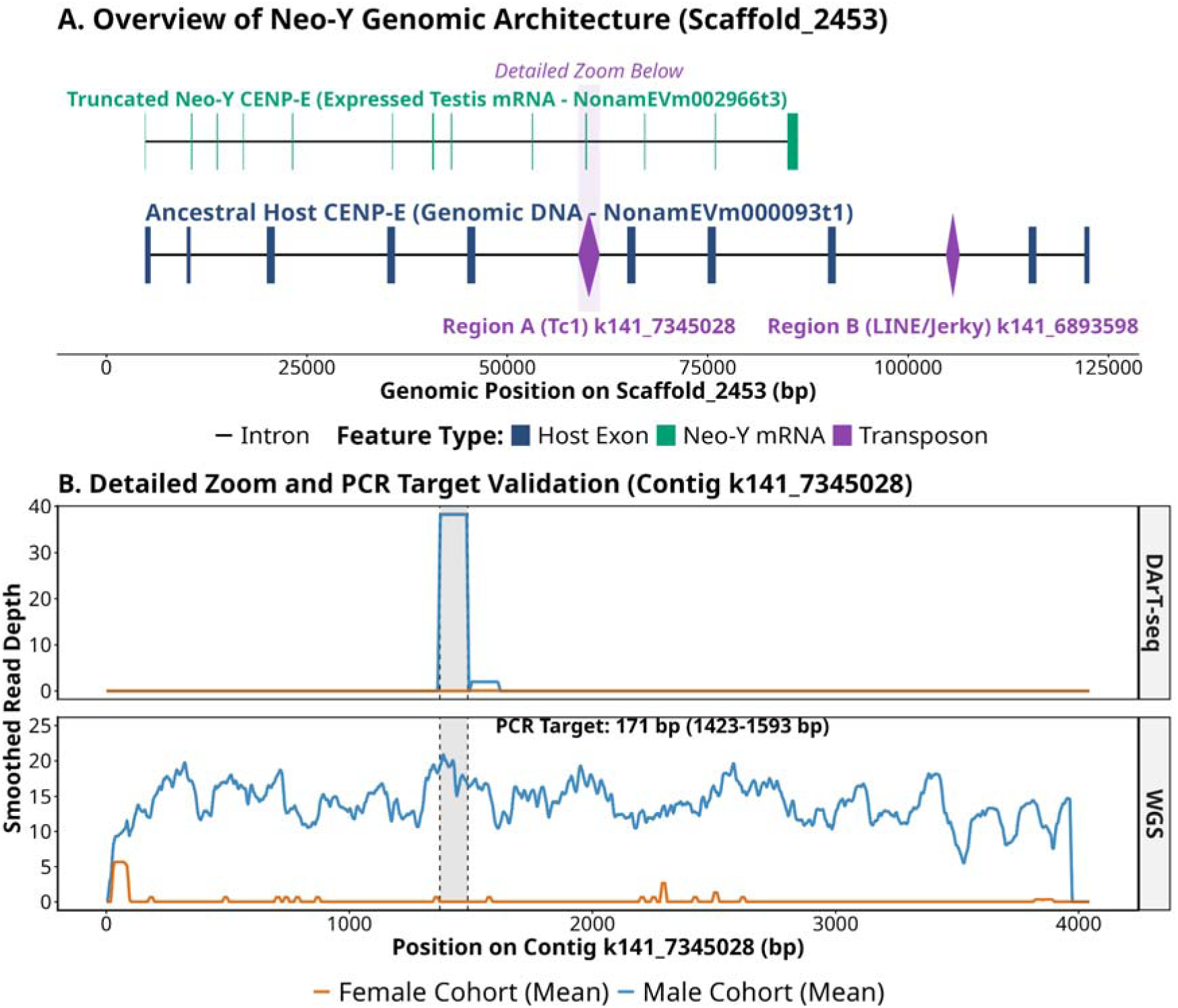
Genomic architecture of the male-specific region highlighting the structural breakpoint used for diagnostic marker design. (A) Structural overview of the male-specific locus on Scaffold_2453, demonstrating the insertion of mobile elements (Region A and Region B) that disrupt the host *CENP-E* sequence. (B) High-resolution read depth validation of the 171 bp diagnostic PCR target spanning the male-exclusive structural breakpoint. DArT-seq and Whole Genome Sequencing demonstrate robust coverage in males (blue) and an absolute absence of reads in the female cohort (orange).

**Figure 5.**
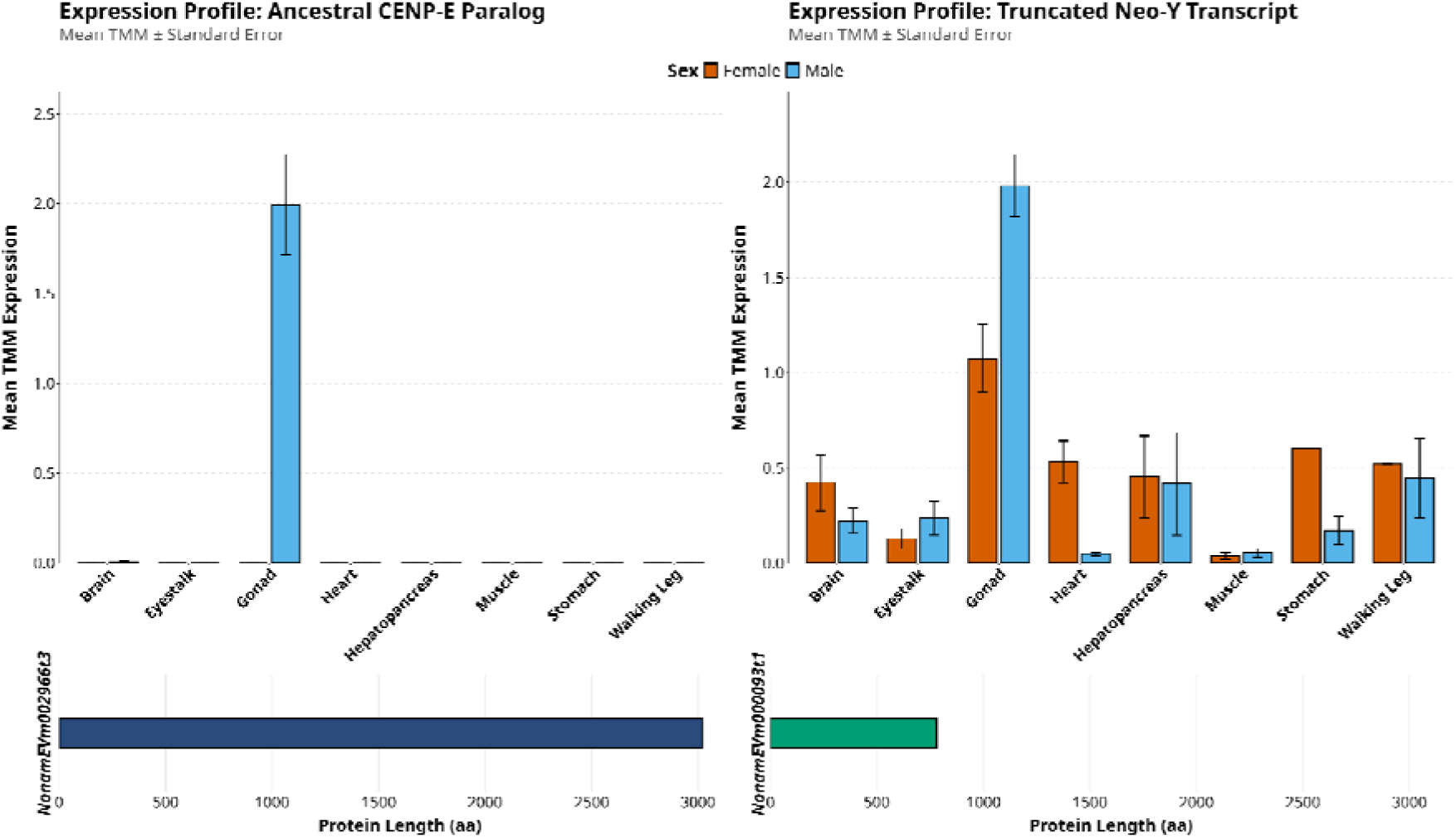
Tissue-specific expression profiles contrasting the somatic ancestral gene and the testis-exclusive male paralog. (A) Transcriptional profile demonstrating broad expression of the ancestral *CENP-E* paralog across multiple somatic and reproductive tissues in both sexes. (B) Expression profile of the newly emerged male *CENP-E* transcript, demonstrating complete silencing across somatic tissues with expression exclusively restricted to the male gonad (testis).

#### 3.2.3. Validation of the diagnostic sex marker

Because the male-exclusive Tc1 insertion acts as a precise structural breakpoint for the neo-Y gene, it provided a flawless target for diagnostic development. To guarantee absolute male-specificity and bypass the false-positive amplification of autosomal Tc1 paralogs in females, we employed a stringent junction-based primer design strategy.

In silico comparative read-depth analysis pinpointed a highly specific 115 bp candidate window (positions 1372–1487) defined by a complete and uniform loss of coverage in the female cohort. Within the individual coverage heatmap, a prominent “white corridor” illustrates the absolute consistency of this absence across all female biological replicates, cementing the region’s diagnostic potential (Supplementary Figure S3).

Capitalizing on this validation, diagnostic primers (SL-SM1) were designed to flank this male-exclusive window, yielding a definitive 171 bp amplicon (Supplementary Table S7.1). Subsequent PCR-based screening across a wild-caught cohort demonstrated the marker’s robust diagnostic power: the 171 bp sequence was consistently amplified in all male SL individuals while remaining entirely undetectable in females (Figure 6A). Blind testing using this marker enabled the flawless genetic identification of 12 male and 12 female samples, achieving 100% phenotypic concordance. The successful co-amplification of the autosomal positive control (SL-PC) across all samples confirmed the quality of the genomic DNA, completely ruling out false negatives in the female cohort and definitively validating SL-SM1 as a strictly male-specific, industry-ready diagnostic tool (Figure 6B).

**Figure 6.**
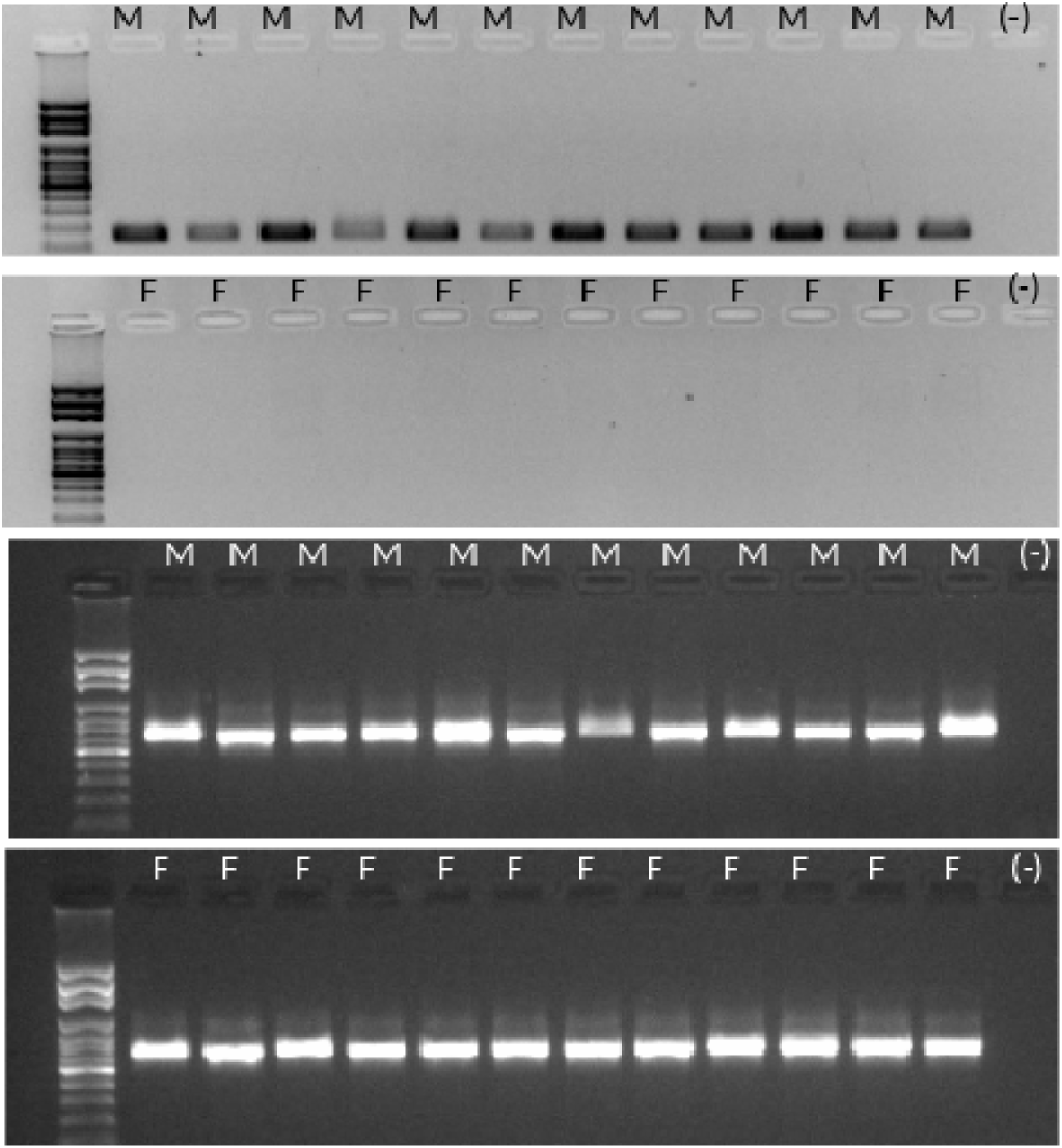
Experimental validation of the 171 bp male-specific diagnostic PCR marker (SL-SM1) demonstrating 100% phenotypic accuracy. (A) PCR amplification using the SL-SM1 marker across 24 wild-caught *T. australiensis* individuals. The 171 bp amplicon is strictly amplified in males (M), whereas no specific marker is amplified in females (F). (B) Co-amplification of the autosomal positive control (SL-PC) confirms high-quality DNA extraction across all 24 samples, ruling out false negatives.

### 3.3. DArT-seq validation of the XX/XY chromosome system

To further validate the sex-determination system at the population level, we analyzed the DArT-seq dataset. A total of 40,602 alleles were generated, of which 4,403 were detected in at least 10 individuals. Among these, 226 alleles displayed a clear sex-biased distribution. Heatmap clustering showed strong segregation of individuals by sex, forming two distinct male and female clades.

Interestingly, 74 alleles were present in the male cohort but were completely absent across all 13 female samples, confirming the presence of strictly male-specific genomic regions (Figure 7A). Subsequently, 37 male-specific sequences were subjected to BLAST analysis against the SL reproductive transcriptome (testis and ovaries). Fourteen of these sequences showed significant matches (*E-values* < 10[^20^), suggesting high sequence similarity and transcriptional relevance. Remarkably, several SNPs displayed read-depth coverage of approximately 50% in testis tissues and 0% in the ovaries. This specific 50% read-depth ratio is highly indicative of heterozygosity mapping against a diploid background, providing strong genomic validation that *T. australiensis* utilizes an XX/XY sex chromosome system characterized by Y-linked regions (Figure 7B).

**Figure 7.**
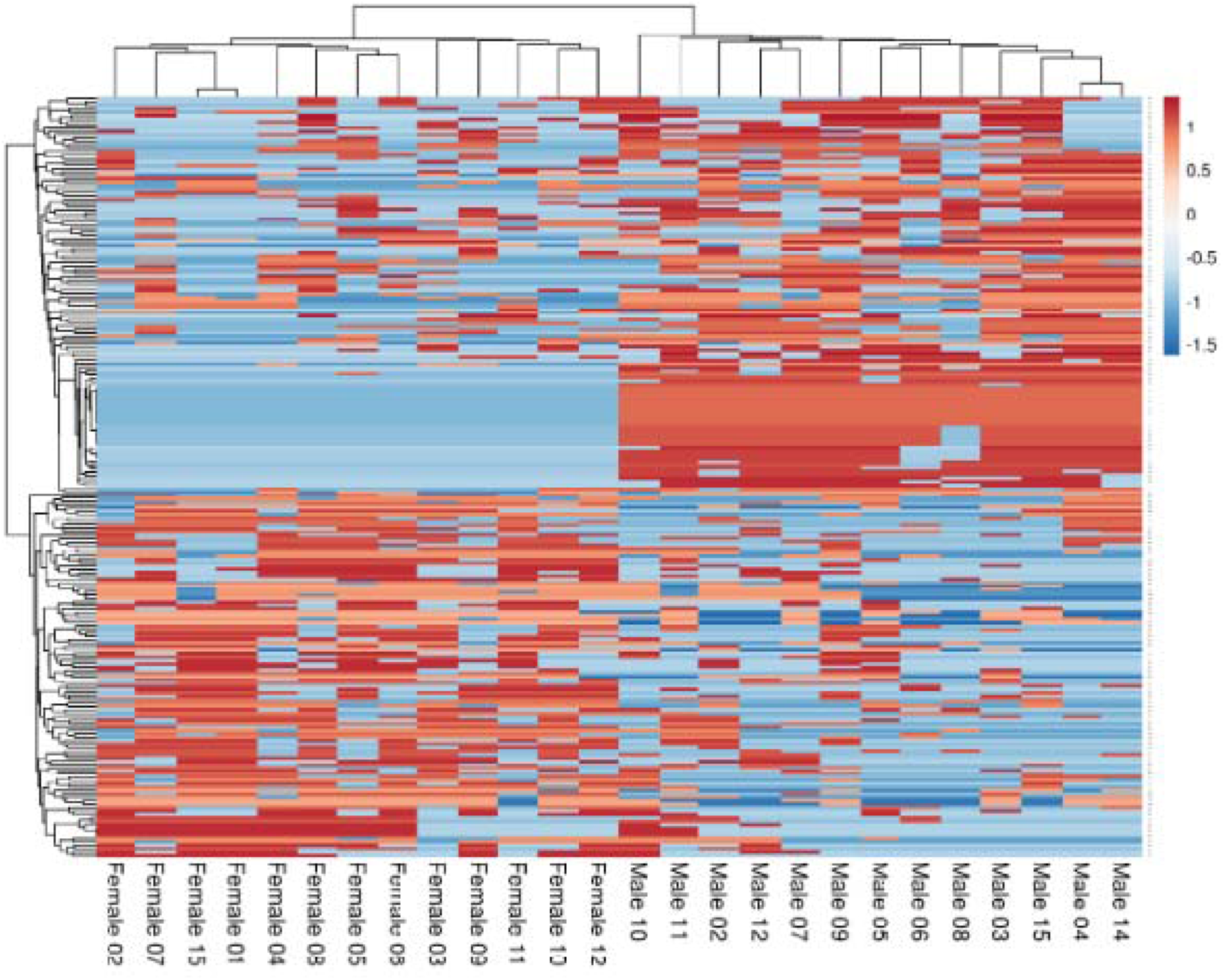
Population-level DArT-seq analysis providing genomic evidence for an XX/XY sex chromosome system. (A) Heatmap clustering of 226 sex-biased DArT-seq alleles showing clear separation of female and male samples; (B) Read alignment and depth profile of male (testis) and female (ovaries) samples showing high and continuous coverage across a male-specific contig, with SNPs displayed around 50% read depth (shown in the red circle), together with their complete absence in females indicates Y-linked genomic regions under an XY sex chromosome system.

## 4. Discussion

*Thenus australiensis* (SL), the most prevalent *Thenus* species in Australia, is a prime candidate for intensive closed-cycle aquaculture due to its brief larval development, accelerated growth, negligible cannibalistic behavior in captivity, and high market value. The development of a draft male genome for SL provides a critical baseline genomic resource. These insights are essential for establishing reliable breeding programs, developing monosex populations, and promoting long-term genetic improvement in sustainable aquaculture practices.

In this study, following rigorous decontamination using the NCBI Foreign Contamination Screen (FCS), the final curated genome size of SL was 0.913 Gbp. This curated assembly represents approximately 73% of our initial k-mer-based genome size estimate of 1.25 Gbp, and remains smaller than its closest phylogenetic relatives, the ornate spiny lobster (*Panulirus ornatus*, 2.65 Gbp) (Ren et al., 2024) and the American lobster (*Homarus americanus*, 2.29 Gbp) (Polinski et al., 2021). This discrepancy is expected and heavily influenced by the limitations of short-read sequencing, which notoriously underrepresents highly repetitive regions, leading to a conservative estimate of the final physical genome size (Berg et al., 2025; Treangen and Salzberg, 2012). While comprehensive annotation of the entire repeatome remains challenging due to fragmentation (Peona et al., 2021; Tørresen et al., 2019), the successfully annotated simple sequence repeats and non-coding RNAs are broadly consistent with other decapod crustaceans (Wei et al., 2025; Yu et al., 2025; Zhang et al., 2019).

Notwithstanding the fragmentation inherent to short-read data, the curated genome exhibits an exceptionally high level of functional completeness, indicated by a BUSCO score of 93.0% against the Arthropoda dataset (Dominguez Del Angel et al., 2018). Utilizing this clean assembly and abundant multi-tissue RNA-seq data, our integrated pipeline predicted a refined set of 30,100 protein-coding genes. This curated gene count aligns closely with the biological average of ∼25,000 genes typically found in other crustaceans (Yuan et al., 2023). The initial assembly prior to decontamination artificially inflated this number, a phenomenon similarly observed in early short-read crustacean assemblies (Veldsman et al., 2021), which was later resolved (Denton et al., 2014). Notably, the number of functionally annotated proteins obtained here (14,937 genes) is highly consistent with the latest chromosome-level assembly of *P. ornatus* (14,884 genes) (Ren et al., 2024). This strongly supports the reliability of our curated gene space, providing a robust genomic foundation for downstream biotechnological applications.

Multiple independent lines of genomic, transcriptomic, and molecular evidence firmly establish that SL utilizes a male-heterogametic (XX/XY) sex-determination system. The structural basis of this system was identified through the discovery of a male-specific *CENP-E* locus on Scaffold_2453. In nascent Y chromosomes, the suppression of recombination often leads to the accumulation of transposable elements and structural decay. The integration of our genomic assembly with multi-tissue transcriptomic profiling captured this degenerative process: a duplicated paralog of the *CENP-E* motor protein has achieved testis-exclusive expression and been subsequently invaded by mobile elements (Class II Tc1 and Class I LINE/Jerky). Crucially, transcription of this neo-Y locus halts prematurely precisely at the Tc1 insertion, severely degrading the protein. This transposon-mediated pseudogenization, paired with its restriction to testis-only expression, is a hallmark of Y-chromosome neofunctionalization and subsequent degeneration (Charlesworth et al., 1994; Steinemann and Steinemann, 1992).

This structural mechanism is perfectly corroborated by the population-level allelic patterns observed in the DArT-seq data. Heatmap analysis identified 74 strictly male-biased alleles present in the male cohort but entirely absent across all female biological replicates. The read-depth coverage of these male-specific SNPs in testis tissue consistently hovered at approximately 50%, while dropping to 0% in ovary tissue. This ∼50% read depth is the exact mathematical signature expected of a heterozygous, single-copy Y-linked locus mapping against a diploid autosomal background, mirroring the evolutionary trajectory seen in other non-model taxa with confirmed XX/XY systems (Lambert et al., 2016).

The most critical and practical outcome of this work is the translation of these genomic discoveries into a highly accurate diagnostic assay. Because the male-exclusive Tc1 insertion acts as a precise structural breakpoint for the neo-Y gene, it provided an ideal target for developing a highly reliable genetic marker. We developed a rapid PCR assay (SL-SM1) that yields a definitive 171 bp amplicon exclusively in males. Subsequent screening across a wild-caught cohort demonstrated 100% accuracy in phenotypic sex identification, with no false positives in females.

For commercial hatchery operations, the implementation of this marker is highly advantageous. DNA can be easily extracted using non-lethal sampling methods, such as taking a minor swimmeret clip from juvenile lobsters. The assay relies on standard PCR equipment, making it cost-effective and accessible for commercial laboratories without the need for advanced sequencing infrastructure. By facilitating early and completely accurate sex identification, this diagnostic tool enables the rapid culling or diversion of slower-growing males. This allows hatcheries to concentrate feed, labor, and valuable tank space entirely on the larger, more profitable female populations, fundamentally optimizing the commercial production of monosex slipper lobsters.

## 5. Conclusion

This study establishes the first heavily curated, functionally complete draft genome for the slipper lobster, *T. australiensis*. While short-read sequencing predictably resulted in a physically fragmented assembly, rigorous decontamination and the integration of multi-tissue RNA-seq data ensured a highly accurate gene space, evidenced by a 93.0% BUSCO completeness score (Dominguez Del Angel et al., 2018) and a robust repertoire of 30,100 predicted protein-coding genes.

Through integrated genomic and transcriptomic analyses, we provide definitive evidence that *T. australiensis* utilizes an XX/XY sex-determination system. The identification of a male-specific *CENP-E* paralog, which has undergone structural decay via transposon invasion, clearly highlights the genomic architecture characterizing the Y-linked region in males.

Crucially, we translated this foundational discovery into a practical, industry-ready diagnostic tool. By targeting a male-exclusive Tc1 insertion, we developed a 171 bp PCR marker that achieved 100% accuracy in phenotypic sex identification. This highly reliable, cost-effective assay eliminates the time required for prolonged progeny testing and provides the critical framework required for early larval sex sorting. Ultimately, these curated genomic resources and the validated diagnostic marker pave the way for the highly profitable commercial production of monosex populations in slipper lobster aquaculture

## Acknowledgments

Funding

This work was supported by the Australian Government with funding from the Australian Research Council (http://www.arc.gov.au/) Industrial Transformation Research Hub (project number IH190100014). The views expressed herein are those of the authors and are not necessarily those of the Australian Government or Australian Research Council.

## Data Availability Statement

Data is contained within the article and Supplementary Materials

## Code availability

In the present study, no custom code was developed. All commands and pipelines used for data processing are detailed comprehensively in the methods section. For software where specific parameters are not explicitly mentioned, we adhered to the default settings as recommended by the software developers. The core code is available at https://github.com/huyha1314/Thenus_australiensis_sex_marker

## Data Records

The raw DArT-Seq and WGS Illumina sequencing data have been deposited in the NCBI Sequence Read Archive (SRA) under BioProject accession PRJNA1394323. The genome-wide shotgun project and final genome assembly have been deposited in DDBJ/ENA/GenBank and are currently undergoing processing. Final accession numbers will be provided prior to publication. To facilitate immediate access for peer review, the complete genome assembly and its corresponding structural annotation information have been made available on Figshare (DOI: 10.6084/m9.figshare.32597610).

