## Supplementary figures and images for "A Draft Male Genome Assembly of the Slipper Lobster *(Thenus australiensis)* Reveals an XY System and a Validated Diagnostic Marker for Monosex Aquaculture"

### FIgure S1

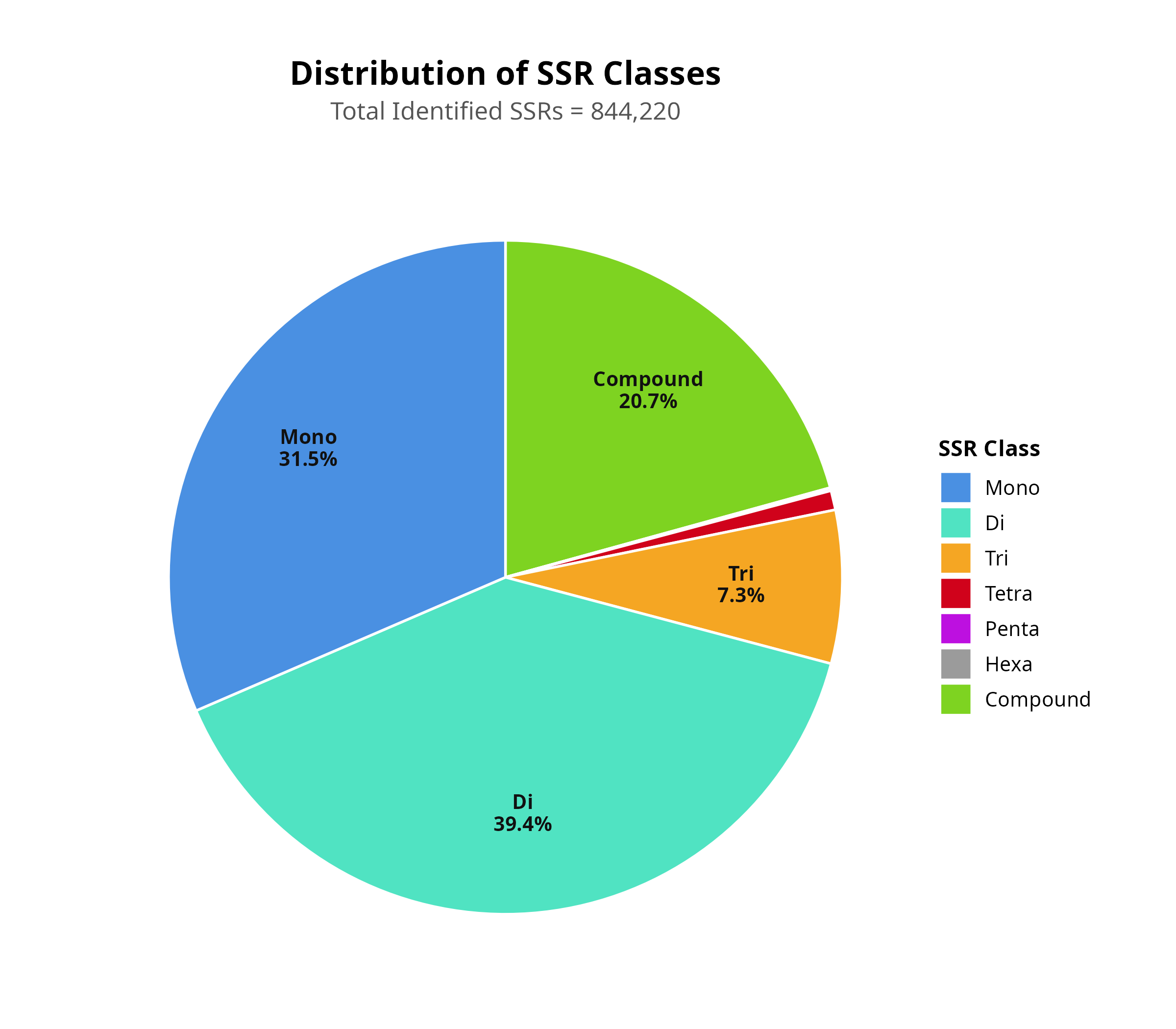

### FIgure S2

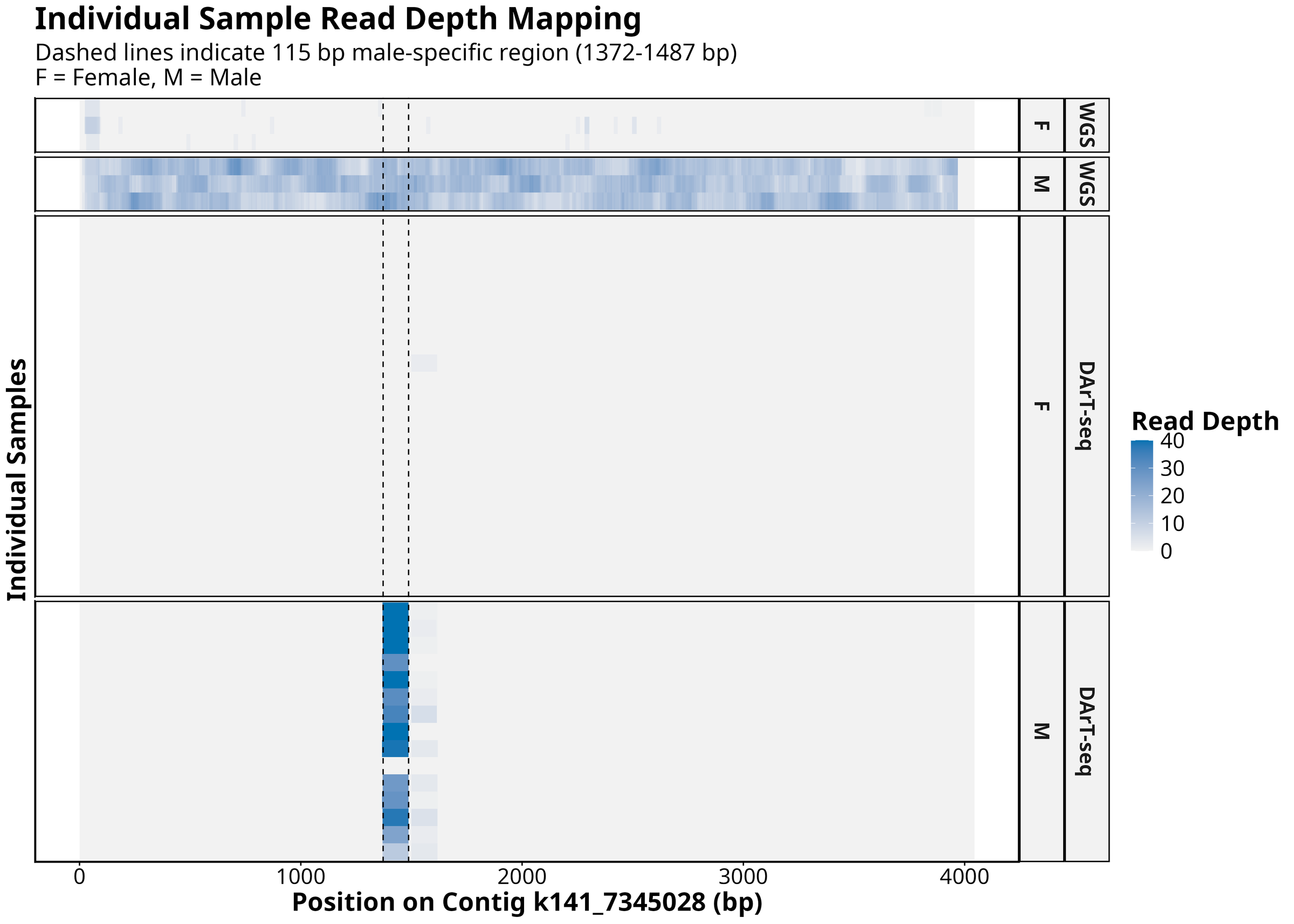

### FIgure S3

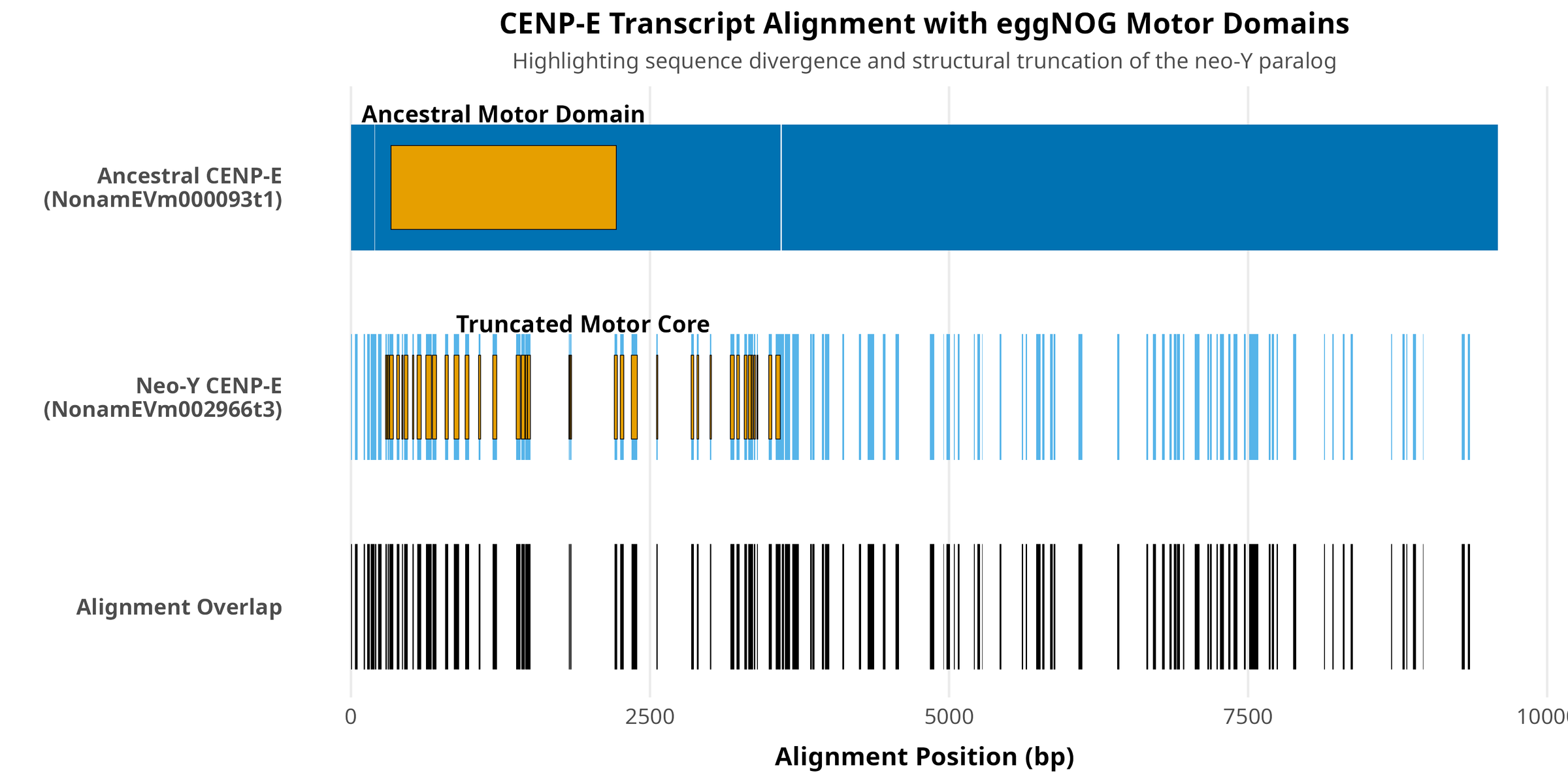
