## Supplementary figure for "A Draft Male Genome Assembly of the Slipper Lobster *(Thenus australiensis)* Reveals an XY System and a Validated Diagnostic Marker for Monosex Aquaculture"

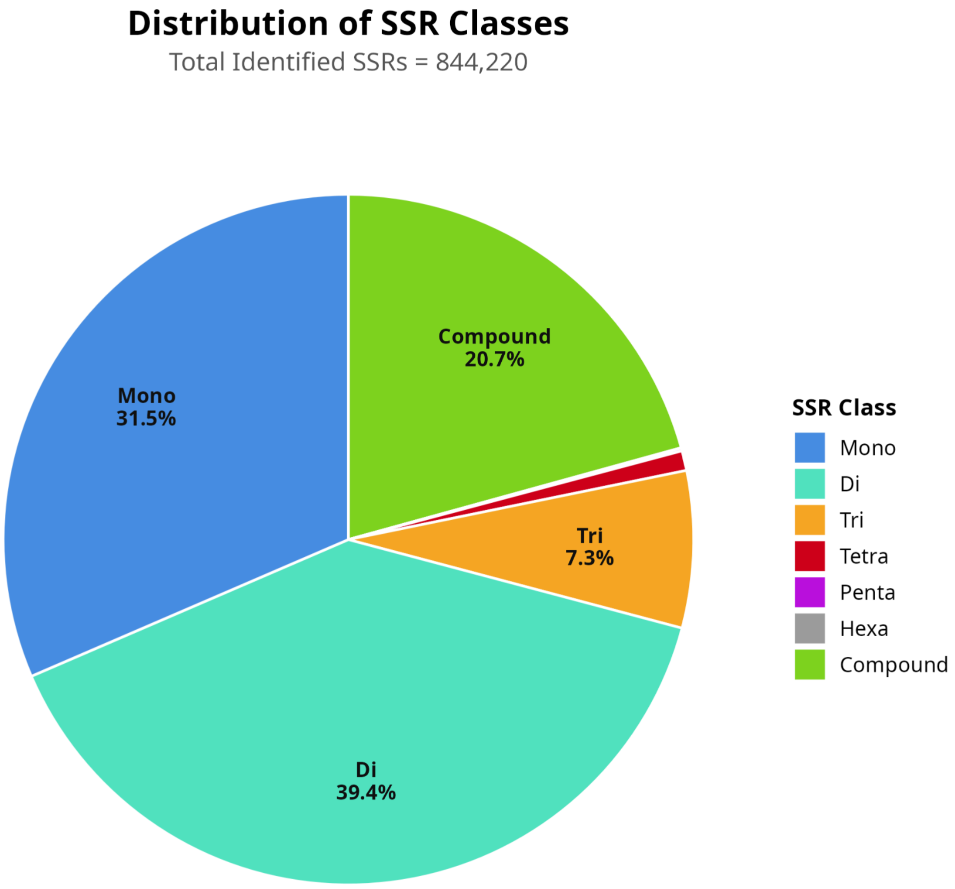


**Supplementary Figure S1. Classification and genomic distribution of repetitive elements in the male *Thenus australiensis* draft genome.** The relative proportions of major repeat classes—including Long Terminal Repeats (LTRs), Long Interspersed Nuclear Elements (LINEs), Short Interspersed Nuclear Elements (SINEs), and DNA transposons—are shown as a percentage of the total annotated repetitive content. Unclassified repeats represent novel or highly divergent sequences unique to the *T. australiensis* genome. Repeat identification and classification were performed using a combined de novo and homology-based annotation pipeline.

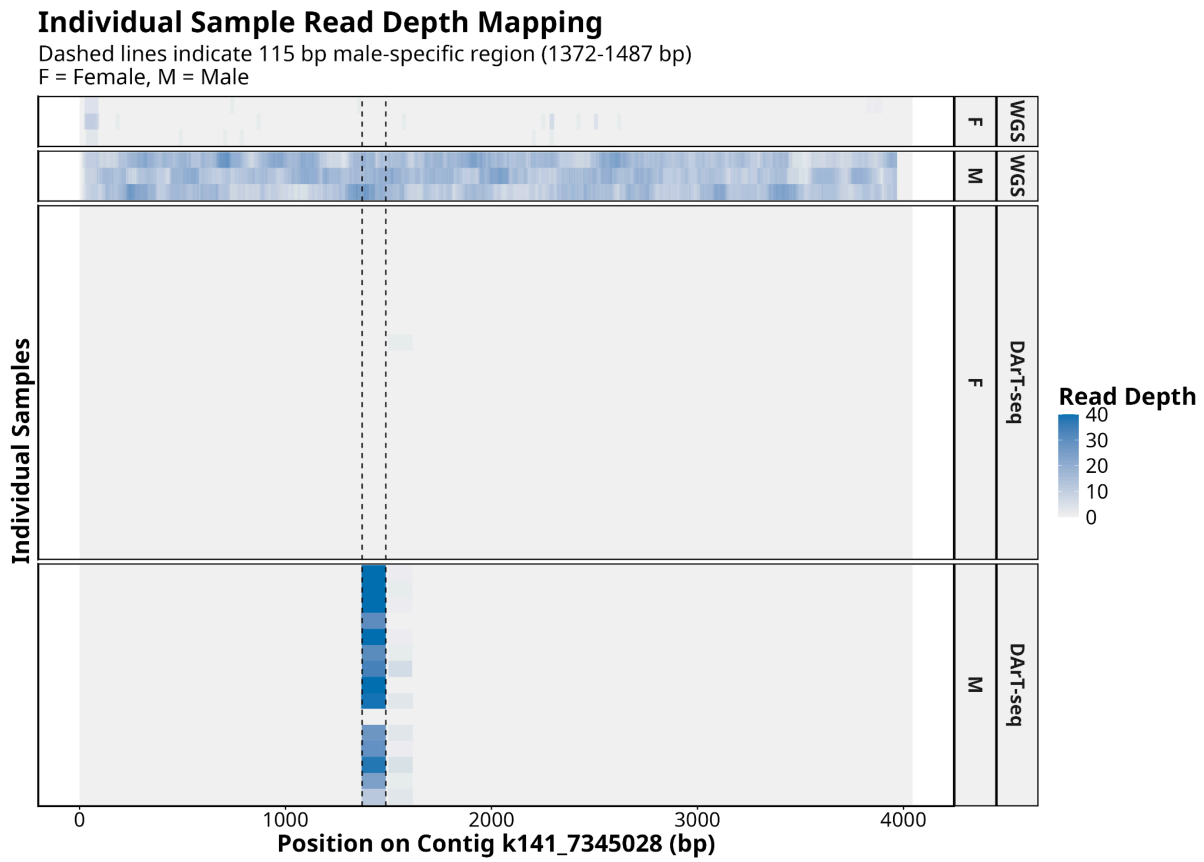


**Supplementary Figure S2. Individual sample read depth validation of the male-specific region on Contig k141_7345028.** Heatmap displaying the sequence coverage across individual female (F) and male (M) samples, evaluated via Whole Genome Sequencing (top panels) and DArT-seq (bottom panels) platforms. The color gradient denotes absolute read depth per sample, ranging from zero (light grey) to high coverage (dark blue). Dashed vertical lines demarcate a strictly male-specific 115 bp diagnostic window (1372–1487 bp). Across both sequencing technologies, this targeted region demonstrates robust, uniform read alignment exclusively in male samples, contrasting with a complete absence of mapped reads in the female cohort, visually confirming the strict sex-linkage of the locus at the individual level.

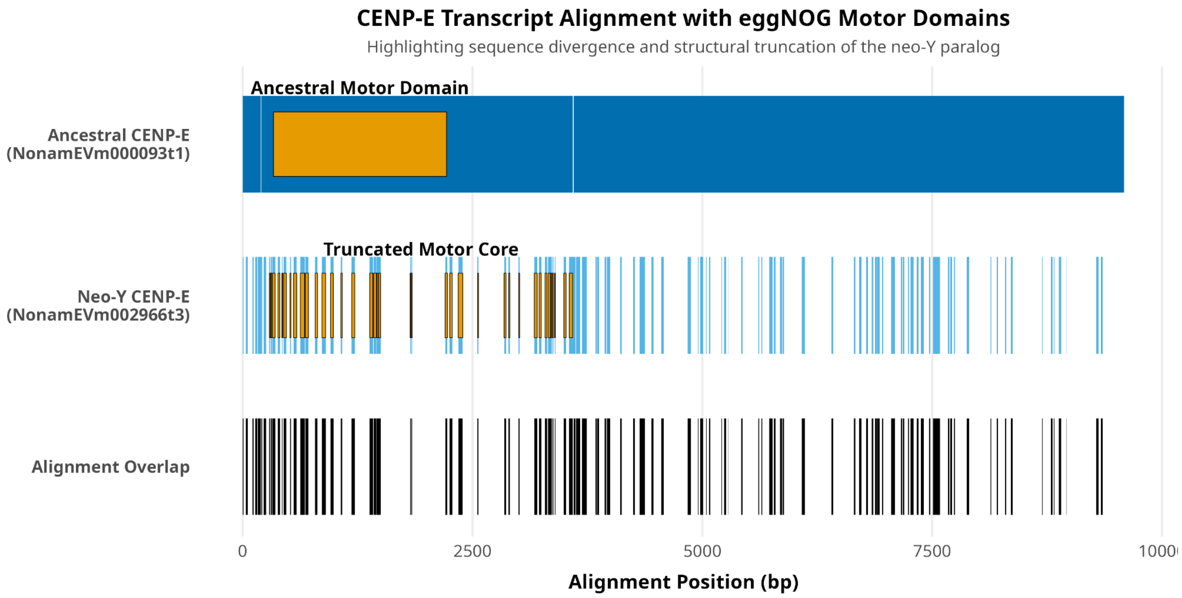


**Figure S3. Structural divergence and domain truncation of the neo-Y *CENP-E* paralog.** Multiple sequence alignment (MSA) mapping of the ancestral autosomal *CENP-E* transcript (top, blue) and the testis-specific neo-Y paralog (middle, pink). Colored blocks indicate regions of mapped sequence coverage, while gaps in the tracks represent insertions/deletions. The bottom track (dark grey) illustrates regions of exact nucleotide overlap between the two transcripts, highlighting the severe structural divergence (60.26% identity; 24.67% coverage). Yellow boxes anchored above each transcript demarcate the kinesin-7 motor domains annotated via eggNOG. Note the premature termination of the neo-Y transcript and the corresponding reduction of its motor domain to the 899 bp bare-minimum catalytic core, contrasting with the full-length 1,883 bp ancestral domain.
